# Larval diapause slows adult epigenetic ageing in an insect model, *Nasonia vitripennis*

**DOI:** 10.1101/2025.05.22.655466

**Authors:** Erin E.B. Foley, Christian L. Thomas, Charalambos P. Kyriacou, Eamonn B. Mallon

## Abstract

Epigenetic clocks based on DNA methylation provide robust biomarkers of biological age, yet the mechanistic basis and functional significance of slowing these clocks remain unclear. Progress has been limited by the lack of short-lived, genetically tractable model organisms with functional DNA methylation systems. The jewel wasp, *Nasonia vitripennis*, offers a unique solution. It combines a functional DNA methylation system with a short lifespan and established tools for experimental manipulation. We previously developed an epigenetic clock in *Nasonia*, but whether this clock reflects plastic, environmentally driven ageing processes was unknown.

Here, we test this directly by experimentally inducing larval diapause, a naturally occurring developmental arrest triggered by environmental cues. Diapause extended median adult lifespan by 36% and significantly slowed the rate of epigenetic ageing. Using whole-genome bisulfite sequencing across multiple adult timepoints, we show that while adults that have passed through diapause as larvae initially emerge epigenetically older, their subsequent epigenetic ageing proceeds 29% more slowly than non-diapaused controls.

Clock CpGs were enriched for gene ontology terms related to conserved nutrient-sensing and developmental pathways, including insulin/IGF signaling and mTOR, supporting the established mechanistic link between development and epigenetic ageing. These findings demonstrate that epigenetic ageing is plastic in *Nasonia* and can be experimentally modulated by early-life environment, establishing this animal model as a tractable system for dissecting the causal mechanisms of epigenetic ageing.

---

Understanding the biology of ageing is a major scientific and societal challenge. Epigenetic clocks, biomarkers based on DNA methylation, have emerged as powerful predictors of biological age and healthspan that can outperform chronological age (1, 2). Yet despite their utility, the mechanistic basis of these clocks and the biological significance of slowing epigenetic ageing remain poorly understood (3).

Progress in this area has been hindered by limitations in the current model organisms. While invertebrates like *Drosophila melanogaster* and *Caenorhabditis elegans* are invaluable for ageing research due to their genetic tractability and short lifespans, they possess only trace amounts of DNA methylation in specific developmental stages or tissues (4, 5), precluding their use for studying the relevance of DNA methylation in ageing.

The jewel wasp, *Nasonia vitripennis*, overcomes this barrier by combining a short lifespan, a well-annotated genome and a functional DNA methylation system (6, 7). We recently established an epigenetic clock in *Nasonia*, making it the first insect model with a methylation-based biomarker of ageing (8). However, it remains unknown whether this clock is plastic and responsive to environmental changes.

Here, we test the plasticity of the *Nasonia* epigenetic clock using diapause, a naturally induced larval developmental arrest triggered by environmental cues. In a number of species, diapause has been associated with altered adult lifespan and with conserved ageing pathways such as insulin/IGF signalling and mTOR, which are thought to mediate the effects of environmental determinants on diapause (9, 10). In *Nasonia*, diapause is also accompanied by broad DNA methylation reprogramming (11).

We show that experiencing larval diapause slows subsequent adult epigenetic ageing in *Nasonia vitripennis*, providing direct evidence that an invertebrate epigenetic clock is responsive to environmental inputs. This establishes *Nasonia* as a powerful new model for dissecting the mechanisms of plastic epigenetic ageing.

## Materials and Methods

Diapause was induced by maintaining *Nasonia vitripennis* mothers at 20°C under a 8:16 h light:dark (LD 8:16) cycle. After ten days, their larval offspring are committed to diapause. Fourth instar diapaused larvae were maintained at 4°C in constant darkness (DD) for three months before adult emergence. All experiments were carried out on virgin males. Post-emergence, diapaused and non-diapaused (control) virgin males were individually housed (25°C, 40% humidity, LD 12:12), fed daily with 20% sucrose solution, and their survival monitored daily.

For whole-genome bisulfite sequencing (WGBS), individuals were sampled at five timepoints post emergence (days 6, 12, 18, 24, 30). Each of the 40 libraries (2 treatments x 5 timepoints x 4 replicates) represented pooled DNA from 10 individuals and included a 1% unmethylated lambda spike-in. Sequencing and bioinformatic processing followed our standard pipeline (8). Agerelated differentially methylated loci (DMLs) were identified (12) and then filtered to retain CpGs strongly correlated with chronological age in control samples for epigenetic clock construction (13). An elastic net regression was trained on age-associated CpG sites to predict chronological age providing an epigenetic clock. Predicted epigenetic age was validated against chronological age using a linear model.

Full methodological details are provided in the Supplementary Information.

## Results and Discussion

Entering diapause as a larva extended adult lifespan (Figure 1A), reducing the hazard of death by approximately 65% compared to controls (Cox Proportional Hazards model; Hazard Ratio [HR] = 0.35, 95% CI: 0.26–0.49; Wald test, *p* = 5 *×* 10^−9^). This was also reflected in the median survival determined by Kaplan-Meier analysis which was 30 days (95% CI: 28–32 days) for the diapause group (n=71), versus 22 days (95% CI: 22–23 days) for controls (n=101).

**Fig. 1.**
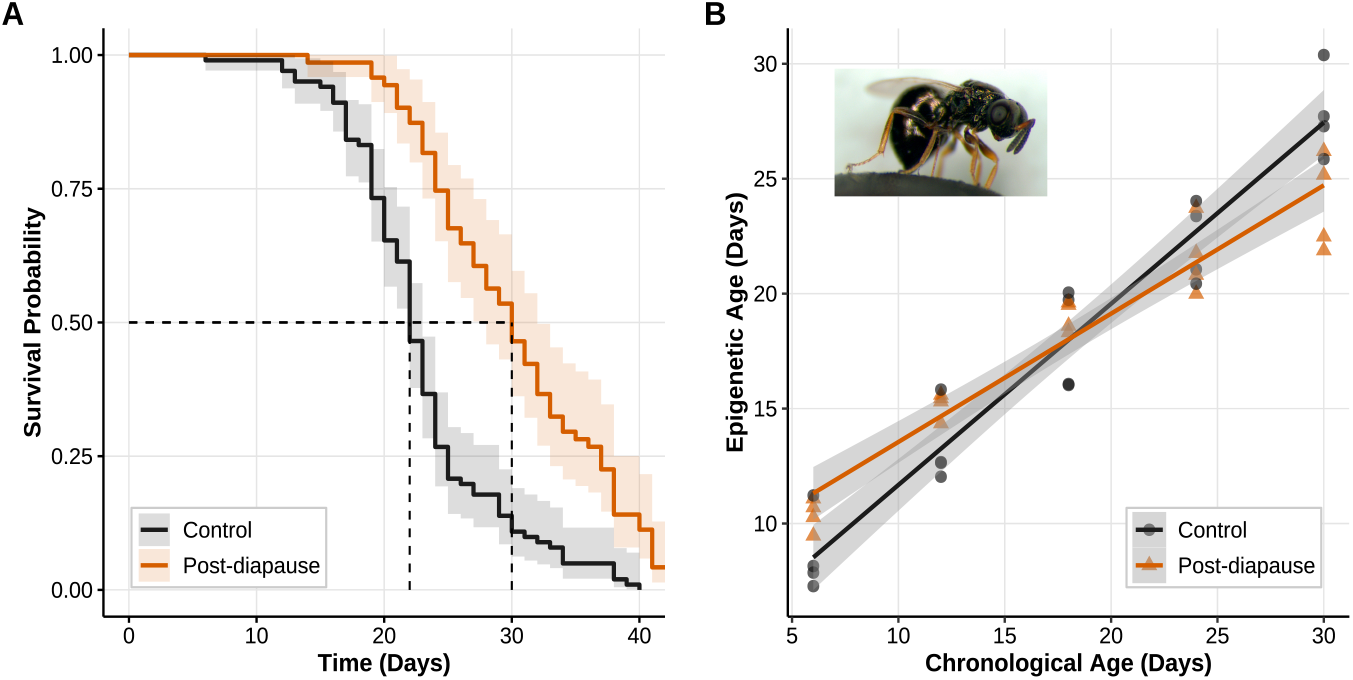
Diapause effect on lifespan and epigenetic ageing. A) Adults diapaused as larvae live longer than non-diapaused conspecifics. Shaded areas represent 95% confidence intervals. Dotted lines represent median survival. B) Diapause slows adult epigenetic ageing. Shaded areas represent 95% confidence intervals. Inset: *Nasonia vitripennis*. Image credit: M.E. Clark public domain

From 715,987 CpG sites classified as methylated (SI Dataset S1), the generalized linear model (12) identified 7,950 CpGs with significant age-related differential methylation (SI Dataset S2). We further prioritized 289 of these sites that were strongly correlated with chronological age (Pearson’s |*r*| ≥ 0.3, uncorrected *p* ≤ 0.05) as input features for an Elastic Net regression model (SI Dataset S3). The final optimised model (mixing parameter *α* = 0.5; regularization *λ* = 1.84207, determined by 10-fold repeated cross-validation [3 repeats] minimizing Root Mean Squared Error [RMSE]) utilized a concise panel of 27 CpGs to estimate epigenetic age (SI Dataset S4).

Our resulting epigenetic clock accurately predicted chronological age in control samples, explaining 91.7% of the variance (cross-validated *R*^2^; RMSE = 2.44 days). Importantly, the clock also performed robustly when applied to diapause samples, accounting for 78.0% of chronological age variance (*R*^2^; RMSE = 3.98 days), demonstrating its potential applicability across distinct physiological conditions.

Curiously, at day 6 post-eclosion, adults that have passed through diapause as larvae were epigenetically older than age-matched controls by an estimated 2.8 days (diapaused: 11.32 days vs. control: 8.53 days, *t* = −3.22, d.f. = 36, *p* = 0.0027, emmeans post hoc). One possible explanation is that epigenetic ageing occurs during the diapause period, albeit at a markedly reduced rate compared to adult ageing. This would suggest that the diapause state, while metabolically suppressed (9), is not entirely inert at the molecular level. Alternatively, the observed overshoot may reflect transient remodelling of DNA methylation during emergence from diapause, as suggested by recent evidence from the silkworm, *Bombyx mori* (14). Distinguishing between these scenarios will require direct measurement of methylation dynamics during diapause itself.

Despite this initial increase in epigenetic age, adults that have passed through diapause as larvae subsequently age epigenetically 29% more slowly than controls (Figure 1B; control slope = 0.78812, diapause slope = 0.55828; linear model interaction of day and treatment: *t* = −3.903, d.f. = 36, *p* = 0.0003). By day 18, both groups converge on an epigenetic age of approximately 18 days. However, by day 30, diapaused individuals are epigenetically 2.7 days younger than controls (24.71 days vs. 27.44 days; `emmeans` post hoc: *t* = 3.152, d.f. = 36, *p* = 0.0033). To evaluate whether this difference could plausibly account for the observed 65% reduction in mortality hazard (HR = 0.35), we considered a simplified Cox model in which hazard increases exponentially with epigenetic age (see Supplementary Information). If the 2.7-day deceleration fully explained the survival benefit of diapause, each additional day of epigenetic age would correspond to a 47.5% increase in mortality hazard. While this may appear large, it is broadly consistent with human and murine studies reporting 5–10% annual increases in mortality per year of epigenetic age acceleration (15, 16). Given *Nasonia*’s short adult lifespan (Figure 1A), a one-day shift in epigenetic age represents a substantial fraction of the life course, and such a daily hazard effect is not implausible. Although we did not model hazard as a function of epigenetic age directly, this calculation supports the view that the clock reflects biologically meaningful variation in ageing rate.

Our findings place diapause-induced epigenetic deceleration within the broader context of early life programming, where environmental cues can reshape long-term molecular and physiological trajectories (17). The rate of change in the epigenome can depend on early life environmental conditions (18). This is not unexpected, as the epigenome is highly plastic during early development. Such plasticity may be adaptive, allowing early life environments to reconfigure the epigenome in ways that enhance future fitness, consistent with the predictive adaptive response hypothesis (19). In this framework, insect larval diapause would serve as a predictive adaptive response: an overwintering strategy that anticipates a more challenging adult environment. Adults that have passed through diapause as larvae may be under selection to survive longer, facilitating reproductive success in harsher post-winter conditions. In *Nasonia*, our data suggest that this is mirrored at the molecular level by a long-term slowing of the epigenetic clock.

Epigenetic clocks across diverse species consistently highlight key developmental gene sets as predictors of biological age (20). The CpGs comprising our *Nasonia* epigenetic clock are significantly enriched for gene ontology terms related to conserved developmental and nutrient-sensing pathways, including mTOR and insulin/IGF signaling (SI Dataset S5). Protein arginine methyltransferase 1 (PRMT1), identified among our clock genes (LOC100122160), links epigenetic ageing to the insulin/IGF signalling (IIS) pathway. PRMT1 methylates the FoxO transcription factor, blocking its phosphorylation by Akt and thereby promoting its nuclear localisation and activity (21). As FoxO mediates key IIS-regulated processes, including diapause and lifespan, PRMT1 may act as an upstream modulator of IIS effects on epigenetic ageing.

Theoretical links between development and ageing have a long history in evolutionary biology (22). However, not all forms of developmental arrest lead to altered adult lifespan. For instance, the dauer stage in *C. elegans* greatly extends lifespan during the arrested state, but does not slow ageing after normal development resumes (23). In contrast, our findings align with recent work in *Drosophila*, where Kang *et al*. (24) demonstrated that delayed development in prothoracicotropic hormone (PTTH)-null mutants extends lifespan and postpones the onset of age-related transcriptional changes. In *Nasonia*, we observe a similar phenomenon, larval diapause, a naturally induced developmental delay, slows the progression of the epigenetic ageing clock. Notably, PTTH suppression has been implicated in diapause induction across diverse insect species (25). Together, these findings point to a conserved endocrine-epigenetic axis through which developmental timing modulates ageing trajectories.

Epigenetic ageing is influenced by inflammation, cell division, metabolic state, and early-life environment (3). With its compact genome, short lifespan, and functional methylation system, *Nasonia* enables experimental dissection of these processes *in vivo* (6). Our findings demonstrate that epigenetic ageing in this model is not only measurable, but developmentally modifiable. This positions *Nasonia* to address a fundamental translational question: can targeted reductions in epigenetic age improve long-term health and resilience? Future studies using CRISPR-dCas9-based targeted methylation editing, either to accelerate or decelerate epigenetic ageing at specific clock loci, could directly test whether methylation dynamics causally influence lifespan and healthspan (26).

## ACKNOWLEDGMENTS

EBF was supported by a BBSRC MIBTP DTP studentships. CT was funded by a Leverhulme Trust award RPG-2020-363. EM was funded by a BBSRC Pioneer Award APP3335. For the purpose of open access, the author has applied a Creative Commons Attribution (CC BY) licence to any Author Accepted Manuscript version arising

